# Magnetic sensitivity of cryptochrome 4a in domesticated quail with migratory origins

**DOI:** 10.64898/2026.02.19.706734

**Authors:** Rabea Bartölke, Kevin B. Henbest, Jessica Schmidt, Takaoki Kasahara, Daniel R. Cubbin, Jamie Gravell, Marco Bassetto, Glen Dautaj, Tommy L. Pitcher, Patrick D. F. Murton, Ghazaleh Saberamoli, Julia J. Forst, Maryam Khazani, Shambhavi Apte, Carl Olavesen, Holly Hayward, Olivier Paré-Labrosse, Lewis M. Antill, Jingjing Xu, P. J. Hore, Christiane R. Timmel, Stuart R. Mackenzie, Henrik Mouritsen

**Affiliations:** AG Neurosensory Sciences/Animal Navigation, Institut für Biologie und Umweltwissenschaften, Carl von Ossietzky Universität Oldenburg, 26129 Oldenburg, Germany; Department of Chemistry, University of Oxford, Chemistry Research Laboratory, Oxford OX1 3TA, U.K.; Department of Chemistry, University of Oxford, Physical and Theoretical Chemistry Laboratory, Oxford OX1 3QZ, U.K.; Department of Biochemistry and Molecular Biology, University of Southern Denmark, 5230 Odense, Denmark; Centre for Advanced Electron Spin Resonance (CAESR), Department of Chemistry, University of Oxford, Oxford OX1 3QR, U.K.; Research Center for Neurosensory Sciences, Carl-von-Ossietzky Universität Oldenburg, 26111 Oldenburg, Germany

## Abstract

Magnetoreception, the ability of animals to sense the Earth’s magnetic field, is a fascinating biological phenomenon. Cryptochromes, in particular cryptochrome 4a (CRY4a), have emerged as potential key players in mediating magnetic sensing in various bird species. Building on an earlier investigation of magnetic field effects on European robin (*Erithacus rubecula*) CRY4a, we focus here on CRY4a from the common/Japanese quail (*Coturnix coturnix/japonica*). Japanese quail is one of the very small number of domesticated bird species whose wild forms are migratory. A detailed spectroscopic study of purified quail CRY4a shows that it has magnetic properties similar to robin CRY4a, suggesting that the quail could be a promising additional experimental model with which to unravel the intricacies of magnetoreception in migratory birds.

## 1. Introduction

The study of magnetoreception has gained significant momentum recently, with an increasing focus on understanding the molecular mechanisms that underlie this extraordinary sensory ability in birds [1-14] and other animals [15-23].

Most studies of bird orientation behaviour in the laboratory have used wild-caught migratory passerines, such as European robins (*Erithacus rubecula*) or Eurasian blackcaps (*Sylvia atricapilla*), which cannot routinely be bred in captivity [24, 25]. The breeding and maintenance challenges in captivity posed by wild-caught migratory passerines preclude gene-editing, which could be an extremely valuable tool in confirming or disproving current magnetoreception hypotheses. In contrast, domesticated birds are available all year round, come with the potential for genetic editing, and require fewer ethics permits. The challenge is that most domesticated birds have been bred for egg or meat production, are not migratory, and do not show the *Zugunruhe* phenotype (migratory restlessness) that is the basis of well-controlled laboratory assays of magnetic orientation [26-35]. However, among domesticated birds, the quail could be a promising exception and thus potentially an additional experimental model for magnetoreception studies. Quails belong to the family *Phasianidae*, which includes chickens, pheasants, partridges, and turkeys, and which diverged early from other avian lineages, making them phylogenetically distinct from passerine birds, [12, 36, 37]. Importantly, their wild forms, the very closely related common quail (*Coturnix coturnix*) and Japanese quail (*C. japonica*), are long-distance, night-migratory birds [38, 39]. Wild quails are suitable for Emlen funnel orientation experiments and show magnetic compass orientation capabilities [40]. The *Zugunruhe* phenotype is attenuated in domestic quail but it can be reacquired by crossing or hybridisation with wild quail [41]. Given that domestic quail, which were originally domesticated from Japanese quail [42], or hybrid quail (common ×Japanese), could potentially become models for magnetoreception research, it is crucial to show that the magnetoreceptor in domesticated quails retains its magnetic sensitivity.

Currently, the leading hypothesis for the mechanism of the magnetic compass sense in birds is based on quantum effects in radical pairs generated by blue-light activation of a cryptochrome protein [1, 2, 20, 21, 43-45]. In most birds, there are three cryptochrome genes, *CRY1, CRY2*, and *CRY4* [12], for which six isoforms have so far been reported: CRY1a, 1b, 2a, 2b, 4a and 4b [4, 46-55]. To function as a magnetic sensor, a cryptochrome must bind the flavin adenine dinucleotide (FAD) cofactor which absorbs the light required for the formation of radical pairs [1, 2, 56]. CRY4a is currently the most promising primary magnetoreceptor: it binds FAD stoichiometrically [57] and its mRNA levels are upregulated during migratory seasons [51]. The splice variant CRY4b, on the other hand, does not seem to be relevant for magnetoreception [58]. Recombinantly expressed and purified European robin and chicken (*Gallus gallus*) CRY4a (*Er*CRY4a and *Gg*CRY4a) are sensitive to external magnetic fields *in vitro* [2, 14]. At present, CRY4a is therefore the most promising primary magnetoreceptor candidate in birds and the only avian cryptochrome in which we can study the radical-pair mechanism in detail.

CRY4a contains a ∼2 nm chain of four tryptophan residues (the Trp-tetrad) running from the FAD cofactor embedded in the protein to its surface [2, 51, 59] (see Figure 1 for the structure of pigeon CRY4a and the sequence numbers of the four tryptophans, labelled Trp_X_H, X ∈ {A, B, C, D}, with A closest to and D furthest from the FAD). Blue-light excitation of the purified wild-type (WT) robin protein triggers a sequence of four rapid electron transfers along the Trp-tetrad to the FAD, forming four consecutive radical pairs (RP_X_ = FAD^•−^ Trp_X_H^•+^) [2, 9]. The same photochemistry takes place in the W_D_F mutant of *Er*CRY4a, in which the terminal tryptophan (WD = TrpDH) has been replaced with a redox-inactive phenylalanine (F), except that the final radical pair is RP_C_ instead of RP_D_ [2]. Although, *in vitro*, the magnetically sensitive radical pair in WT *Er*CRY4a is RP_D_, there is a possibility that RP_C_ and RP_D_ interconvert rapidly *in vivo*, forming a “composite” radical pair with weighted average properties of its two components [2, 3]. Compared to RP_D_ alone, potential advantages of such an entity are an enhanced magnetic sensitivity (RP_C_ in W_D_F shows larger magnetic field effects than RP_D_ in the WT protein) and more efficient magnetic signalling [2, 3, 14]. Comparison of WT and W_D_F CRY4a may therefore shed light on this hypothesis.

**Figure 1.**
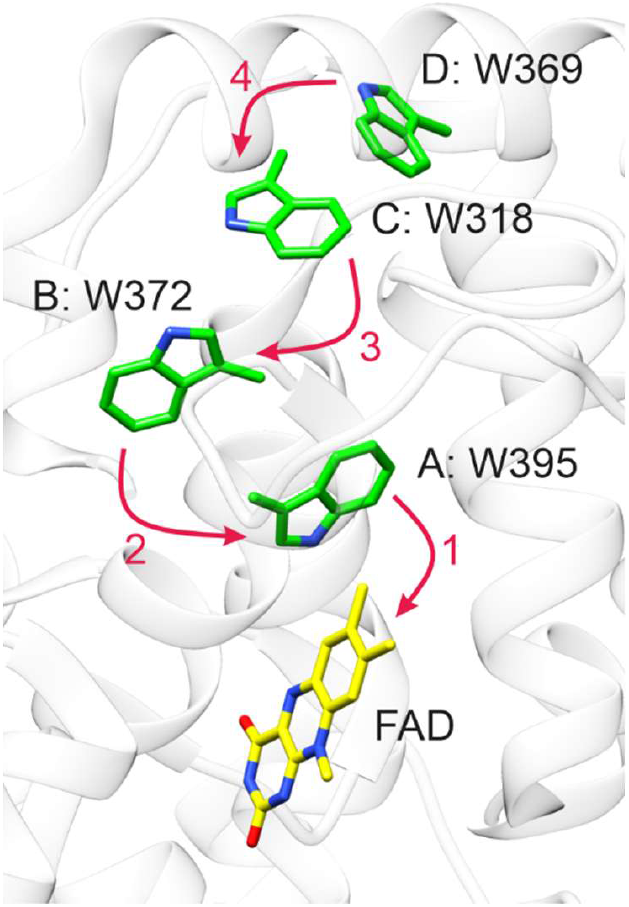
Structure of the electron transfer chain in pigeon *Columba livia (Cl*) CRY4a (PDB: 6PU0 [59]). The arrows indicate the four sequential electron transfers along the tetrad of tryptophan residues to the FAD chromophore. The sequence numbers are common to *Cj*CRY4a, *Er*CRY4a, and *Gg*CRY4a. Only the isoalloxazine part of the FAD is shown.

In this study, we combine complementary forms of spectroscopy to probe light-induced radical pairs in quail *Coturnix japonica* (*Cj*) CRY4a and their sensitivity to magnetic fields. Continuous-wave and pulsed electron paramagnetic resonance (EPR) reveal the distances between the flavin and tryptophan radicals and the internal magnetic (hyperfine) interactions that are essential for the magnetic sensitivity of the radical pairs. Transient absorption (400-800 nm) captures nanosecond-to-microsecond kinetics of flavin excitation, semiquinone formation, and subtle field-dependent modulations in the lifetimes of the radicals. Cavity ring-down spectroscopy and broadband cavity-enhanced absorption spectroscopy boost the optical detection sensitivity, enabling precise quantification of magnetic-field-induced absorbance changes across multiple radical absorption bands. Finally, confocal fluorescence microscopy tracks the time-dependent change in fluorescence intensity in response to an external magnetic field. A mutant with W_D_ replaced by a phenylalanine residue was also created and analysed: preventing RP_D_ formation enabled the investigation of RP_C_ in more detail.

This study was motivated by the potential benefits of studying magnetoreception in a domesticated bird, the sequence similarity of the robin and quail proteins, and the possibility that RP_C_ and RP_D_ could together be the source of magnetic sensing *in vivo*. We report here an investigation of the magnetic responses of purified WT quail CRY4a and its W_D_F mutant, and compare those results to previous studies of WT and W_D_F CRY4a from robin and chicken [2, 7, 14].

## 2. Methods

### 2.1. Protein expression and purification

Quail spleen and brain tissues were obtained from captive-bred quails raised in the animal care facility of the University of Oldenburg. The genomic DNA was extracted from the spleen, and the mitochondrial DNA control region was amplified using the primers PHDL and PH-H521 [60]. Total RNA was isolated from the brain using TRIzol Reagent (Thermo Fisher, Waltham, MA, USA) according to the manufacturer’s instructions. 1 µg total RNA was reverse-transcribed using SuperScript III Reverse Transcriptase (Thermo Fisher, Waltham, MA, USA). *CRY4* primers were designed based on the predicted sequence of *C. japonica CRY1*-like (XM_015884930.2, still annotated in the NCBI database as *CRY1*-like, although it is in fact *CRY4*) and cDNA was amplified using forward primer 5’ATACGACTAGTATGAGGCACCACACC3’ and reverse primer 5’ATACGAAGCTTCTAGGCTTGCTCTGTCATCC3’ and cloned into the pCold vector using *Spe*I and *Hind*III restriction sites, resulting in an N-terminal deca-His-tagged fusion protein. The site-specific W369F mutation was introduced using the Q5 Site-Directed Mutagenesis Kit (New England Lab, Ipswich, MA, USA) and forward primer 5’GGGGGATCTTTTTATCAGCTGGGAAG3’ and reverse primer 5’CTGGTCAGGAAGCAGGCG3’. Protein expression and purification has been described in detail in Ref. [2] with modifications described in Ref. [14]. Isopropyl-D-1-thiogalactopyranoside (IPTG) was added for induction at a final concentration of 10 µM. All resultant purified protein samples were concentrated to 5-6 mg/mL and checked for FAD binding using a Cary 60 UV-Vis spectrophotometer (Agilent Technologies, Santa Clara, CA, USA) (Supplementary Figure S2). Samples were snap frozen in storage solution (20 mM Tris, pH 8.0, 215 mM NaCl, 20% glycerol, 10 mM 2-mercaptoethanol and 0.6 M trehalose) and shipped on dry ice from the production site in Oldenburg to Oxford, where they were stored at −80 °C until they were prepared for subsequent measurements.

### 2.2. Identification of CRY4 variants in wild quails

Whole genome shotgun sequencing data from wild quails (*C. coturnix* and *C. japonica*) were found in the Sequence Read Archive [61] (BioProject accession no.: PRJNA339911, PRJNA730394, PRJNA1069095, PRJNA1153307). The coding region of *CjCRY4* (XM_015884930.2) was split into each exon, and these were used as queries for the BLASTN search. The resulting reads were displayed in the NCBI Multiple Sequence Alignment Viewer to identify variants and determine whether they were heterozygous or homozygous. Only data from individual birds with coverage of 5 or more in all regions are shown in Table 1. Possible effects of missense variants were predicted using PolyPhen-2 (http://genetics.bwh.harvard.edu/pph2/), one of the most widely used variant effect predictors [62]. PolyPhen-2 was trained on mutations found in humans and their functional effects, but it also takes into account evolutionary conservation and steric structure, including non-human organisms, to make predictions. To predict the effects of *Cj*CRY4 variation, PolyPhen-2 used an alignment of (at most) 75 amino acid sequences of vertebrate CRY1, CRY2, and CRY4 proteins, as well as the crystal structures of pigeon CRY4 (Protein Data Bank: 6PU0) and mouse CRY1 (7D19).

**Table 1.**
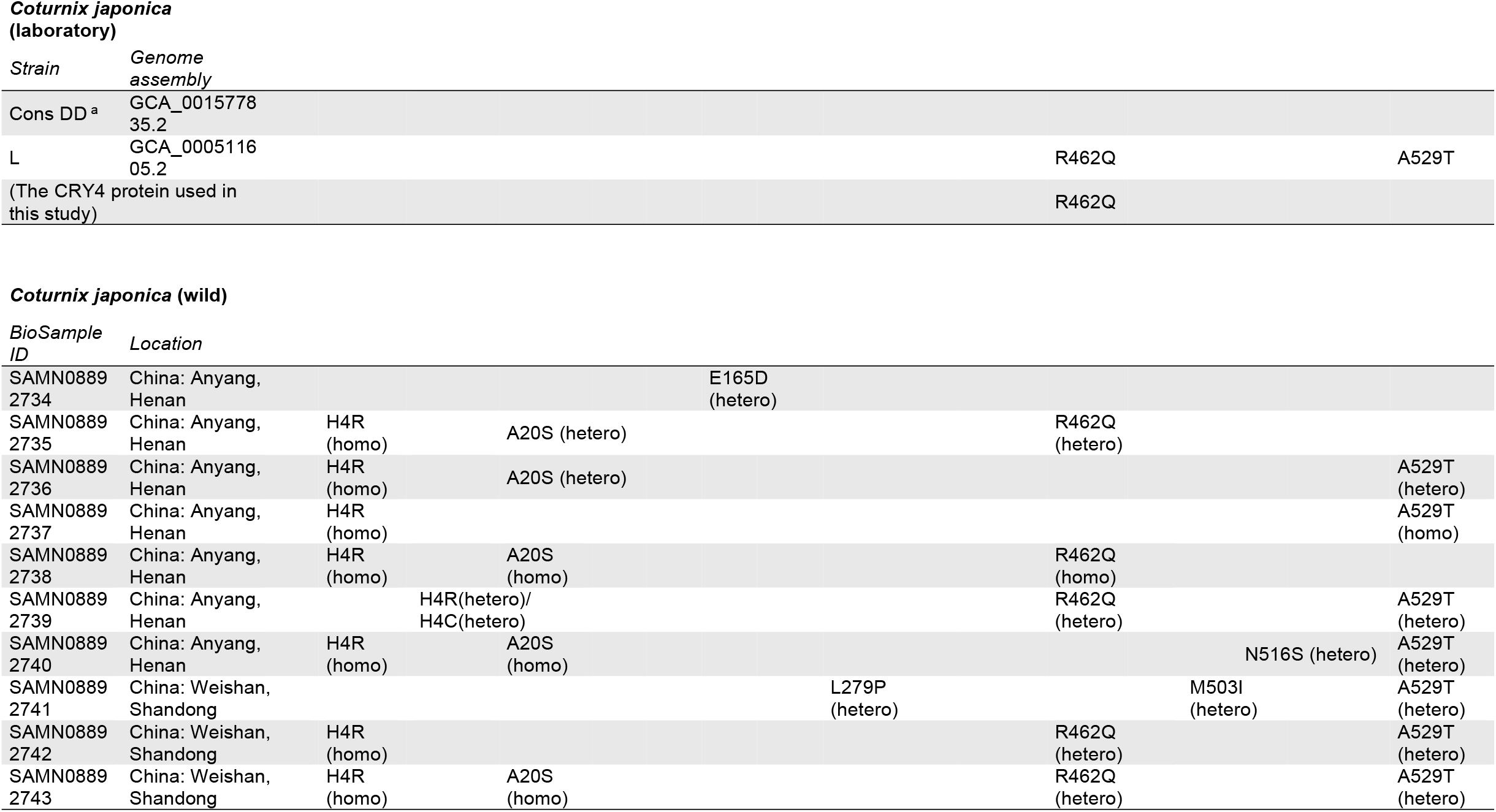
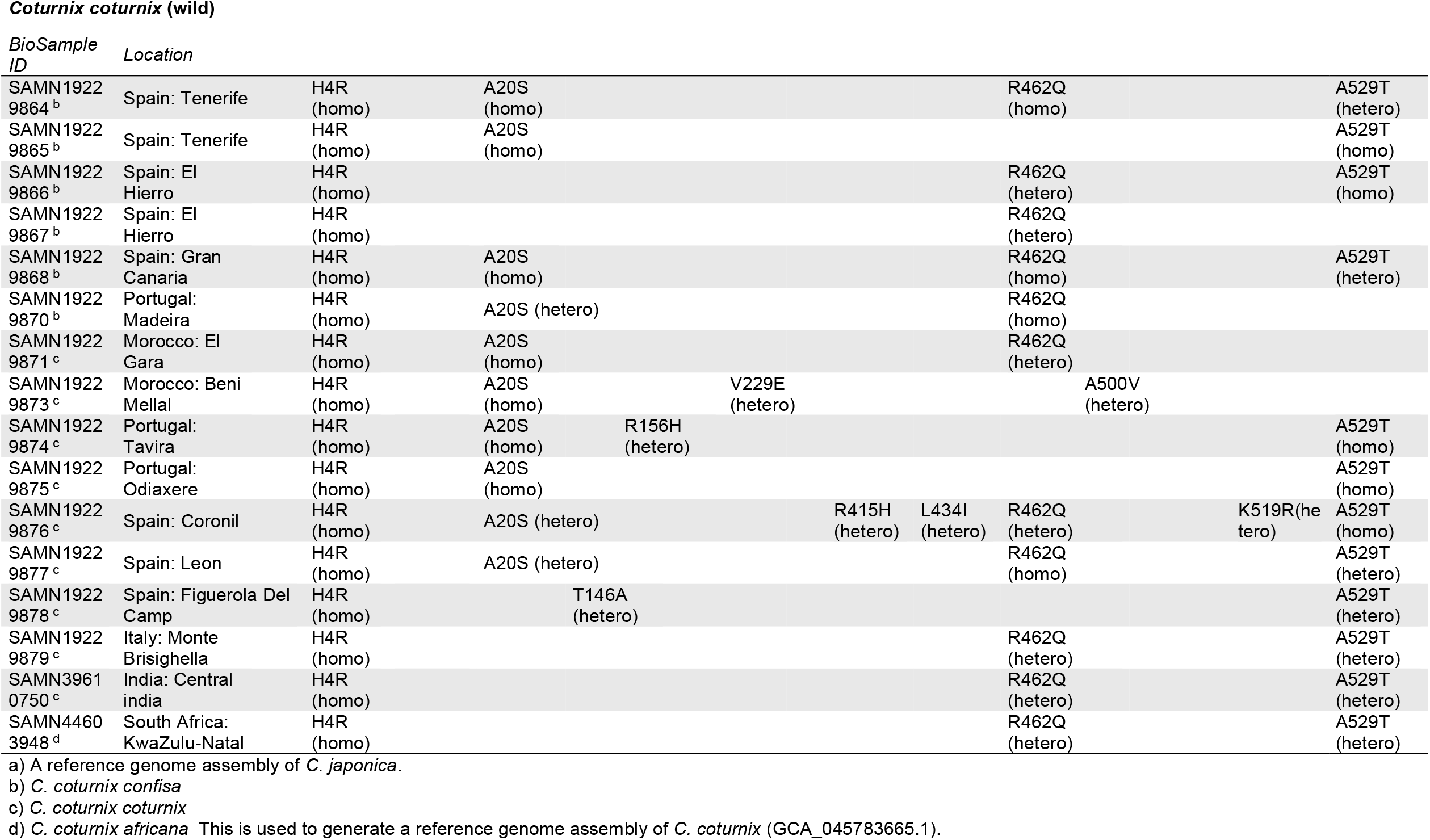
Missense variants in CRY4a found in the quail genome assemblies and wild quail.

### 2.3. Electron paramagnetic resonance spectroscopy

X-band time-resolved EPR (at 274 K) and Q-band out-of-phase electron spin echo envelope modulation (OOP-ESEEM, at 80 K) measurements were performed and analysed as described in Ref. [14]. A Bruker ELEXSYS E580 spectrometer was used in conjunction with either a Bruker MD5 resonator (time-resolved EPR) or a Bruker D2 resonator (OOP-ESEEM). All measurements were made on samples to which 5 mM K_3_Fe(CN)_6_ had been added to promote reoxidation of the flavin. (Addition of K_3_Fe(CN)_6_ allowed faster data collection without affecting the appearance of the spectra). The OOP-ESEEM data, *S*(*τ*), were fit from 200 ns onwards to the expression:

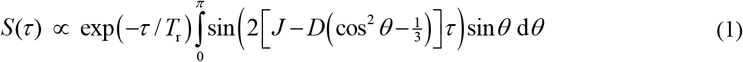

>to extract the dipolar (*D*) and exchange (*J*) couplings and the relaxation time *T*_r_ [63]. From *D*, the inter-radical separation, *r*, was determined, within the point-dipole approximation, as

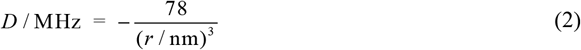

### 2.4. Transient absorption spectroscopy

Transient absorption (TA) spectroscopy was used to follow the first 1 μs of radical formation. An Ultrafast Systems EOS spectrometer was employed as described in Ref. [14]. The 450 nm pump pulse (30 μJ, 15 ps, 1 kHz repetition rate) was provided by a Nd:YAG-pumped optical parametric generator (Ekspla PL2210 and PG403) and the broadband (350-900 nm) probe pulse by a 2 kHz supercontinuum source (1 ns pulse width, Leukos Lasers). Spectra were accumulated with pump-probe delays up to 1 μs. Wavelength-resolved kinetic TA data, Δ*A*(*t*), were fit to a (simplified) biexponential decay plus a constant offset to provide an estimate of the radical lifetimes (see Supporting Information).

Magnetic field effects (MFE) were determined by acquiring Δ*A* data in the presence and absence of a *B* = 25 mT magnetic field:

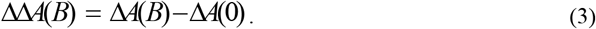

>Significant averaging was required to measure MFEs using TA and our protocols include extensive intervals without irradiation in an attempt to minimise the effects of photodegradation. Nevertheless, for the WT protein it was necessary to add 1 mM K_3_Fe(CN)_6_ as a flavin reoxidant. In the case of the W_D_ mutant, by contrast, the O_2_ in air-equilibrated solutions proved sufficient.

### 2.5. Cavity ring-down spectroscopy

Optical cavities offer significant enhancements in the sensitivity of spectroscopic detection. Cavity ring-down spectroscopy (CRDS) involves monitoring the real-time decay in light intensity oscillating within an optical cavity following injection of a pulse of light. More commonly employed in the gas-phase, we have developed several variants for use in the condensed phase, especially as the probe in pump-probe MFE measurements [7, 64]. Full details of the variant applied here are given in Ref. [7]. The sample was contained within a 1 mm optical path length sample cell with high quality optical windows placed in the middle of a stable optical cavity formed by two highly reflective (*R* > 0.998, 400 nm ≤ *λ* ≤ 800 nm) broadband coated plano-convex mirrors. Photoexcitation at 450 nm was provided by a 300 μJ, 8 ns pulse from a Nd:YAG-pumped dye laser. The probe pulse, predominantly at 532 nm for the measurements included here, was the output of an optical parametric oscillator (Ekspla) injected into the cavity through the front cavity mirror. The per-pass absorbance, *A*, was calculated from the characteristic ring-down time, *τ*, of light circulating within the cavity. Measured via the fraction of light escaping at the back mirror, the absorbance *A*= *L*/(2.303*cτ*), in which *L* is the length of the cavity, *c* the speed of light in air and the factor 2.303 (= ln(10)) converts common (decadic) to natural logarithms. Photoinduced absorbance changes at a pump-probe delay time *t*, Δ*A*(*t*), were determined from the change in *τ* arising from the pump pulse as

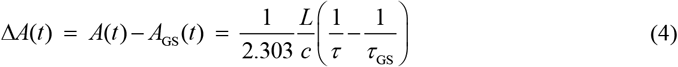

>where *A*_GS_ and *τ*_GS_ (GS = ground state) were measured in the absence of the 450 nm pump pulse.

MFEs were determined from the difference in Δ*A*(*t*) measurements in the presence and absence of an externally applied magnetic field, ΔΔ*A*(*t,B*) = Δ*A*(*t,B*)−Δ*A*(*t*,0) . All CRDS measurements were recorded on samples containing 5 mM K_3_Fe(CN)_6_ to promote flavin reoxidation.

### 2.6. Broadband cavity-enhanced absorption spectroscopy

Another optical cavity technique, broadband cavity-enhanced absorption spectroscopy (BBCEAS) provides continuous broadband spectral resolution [14, 65] from 450 to 800 nm albeit at the cost of time resolution. The details of the BBCEAS implementation used here are given in Ref. [14]. The sample (180 μL, 27 μM, 0.3 optical density) was cooled to 5 °C within a 1 mm path-length optical cell in the centre of an optical cavity and irradiated continuously using a 450 nm diode laser aligned at a small incidence angle to the optical axis. The probe light came from a supercontinuum light source (NKT SuperK Extreme). Although this provided 700 ps pulses at 78 MHz, the time between pulses (13 ns) was much shorter than the characteristic ring-down time of the cavity and the probe was treated as pseudo-continuous. The light exiting the cavity was dispersed in a monochromator (Andor Shamrock) and recorded on a CCD array (Andor Newton). The cavity transmission was converted to absorbance *A*(*t, λ*) as described in Ref. [14] providing photoinduced absorbance changes Δ*A*(*t, λ*) and magnetic field effects, ΔΔ*A*(*t, λ, B*), by analogy with TA and CRDS.

### 2.7. Fluorescence microscopy

An alternative approach to MFE detection in cryptochromes is to use the small fluorescence quantum yield [66, 67] of the flavin chromophore. For this purpose, and reflecting the small sample volumes available, we have developed confocal microscopy for MFE detection, both in bulk solution and for single crystals [14, 68]. A Leica SP8 confocal microscope was adapted to include iron-core solenoid coils for MFE measurements. The microscope works in raster mode with a 100 μm × 100 μm sample volume of protein solutions excited at 448 nm (70 μW) via a 63× oil objective. The total ^1^FAD* fluorescence (475-650 nm) was detected as a 17 mT field was repeatedly switched on and off. MFEs were quantified as differences in the fluorescence intensity, *I*_F_ (which, under constant illumination, reflects the ground state FAD concentration):

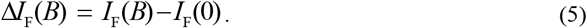

>Sample re-oxidation was achieved, as in Ref. [14], using a custom-built sample holder fitted with an inlet capable of delivering humidified oxygen to the sample. Details of the data analysis, including background subtraction procedure are given in Refs. [14, 68].

## 3. Results

### 3.1. Quail CRY4 sequence

A BLAST search on the NCBI database [69] found a protein sequence (XP_015740416.1) for the Japanese quail that showed high homology to *Er*CRY4a. Although it is annotated as CRY1-like, it is 88% identical to *Er*CRY4a and 96% identical to chicken (*Gg*) CRY4a. Next, using this sequence information, we obtained the cDNA sequence encoding quail CRY4a from brain tissue, which was sampled from quail raised at the University of Oldenburg. Mitochondrial DNA analysis showed that the quail cohort had the most common haplotype detected in wild *C. japonica* and domestic *C. japonica* [70]. The quail CRY4a sequence we obtained differs by one residue (R462Q) from XP_015740416.1; however, this is a natural variant that is found at high frequency in wild *C. coturnix* and *C. japonica* (Table 1). When we studied the CRY4a sequences from wild quails (*C. coturnix* and *C. japonicus*), it was clear that the amino acid sequences resulting from the *CRY4* genes of the two species are identical except maybe at position 4 and some small differences in allele frequencies in other polymorphic amino acid positions, i.e. in amino acid positions where different amino acids occur naturally in the wild populations. Thus, based on the CRY4a sequences alone, it is not obvious that common and Japanese quails are different species (they were once considered subspecies of a single species). The only amino acid location in which the sequence we used in the present study differs from most common quails is remote from the electron transfer chain (position 4), where our sequence had a histidine (H) and most wild common and Japanese quails have an arginine (R). In conclusion, the quail CRY4a protein used in this study is >99% identical to the CRY4a sequence in both the *C. japonica* and *C. coturnix*. As the sequence variant we used is slightly more frequent in *C. japonica* than in *C. coturnix*, we hereafter refer to it as *Cj*CRY4a. The four missense variants (H4R, A20S, R462Q, and A529T) that were recurrently found in both wild *C. coturnix* and *C. japonica* quails were all classified as benign by PolyPhen-2 variant effect predictor [62] (Table 2).

**Table 2.** Polyphen-2 scores of recurrent missense variants in quail CRY4a.

| Variant | Classifier model |  |
| --- | --- | --- |
|  | HumDiv | HumVar |
| H4R | 0.000 (benign) | 0.001 (benign) |
| A20S | 0.020 (benign) | 0.075 (benign) |
| R462Q | 0.012 (benign) | 0.011 (benign) |
| A529T | 0.026 (benign) | 0.011 (benign) |

Wild-type *Cj*CRY4a and its W_D_F mutant carrying W369F were heterologously expressed in *E. coli* and purified to homogeneity for a series of spectroscopic analyses.

### 3.2. Electron paramagnetic resonance spectroscopy

Figures 2a,b show time-resolved 9.637 GHz (X-band) EPR spectra of purified WT and W_D_F *Cj*CRY4a recorded at 274 K, 0.5-1.0 μs after a 5 ns, 450 nm laser pulse. Under these conditions, molecular tumbling is sufficiently slow that the spectra are ensemble averages over an effectively static isotropic distribution of molecular orientations. As expected for a pair of radicals formed simultaneously by light-induced electron transfer, the spectra are spin-polarized: mostly emissive at low field and mostly absorptive at high field [71]. Essentially identical to the corresponding data for the WT and W_D_F forms of *Er*CRY4a, *Gg*CRY4a, and *Cl*CRY4a [2, 14, 54], the *Cj*CRY4a spectra are consistent with the formation of [FAD^?−^ TrpH^•+^] radical pairs in a spin-correlated singlet state. The spectral shapes reflect a combination of *g*-tensor anisotropy, the electron-electron dipolar and exchange interactions, and hyperfine interactions [71-73]. As in the case of *Er*CRY4a, *Gg*CRY4a, and *Cl*CRY4a, the differences between the two *Cj*CRY4a spectra suggest that the radical pair is RP_D_ in the WT protein and RP_C_ in the W_D_F mutant which lacks the fourth tryptophan (W_D_) [2, 14, 54]. The broader appearance of the W_D_F spectrum arises from the larger dipolar coupling (shorter radical-radical distance between FAD^•−^ and Trp_C_H^•+^ in the W_D_F mutant than between FAD^•−^ and Trp_D_H^•+^ in the WT) and the different orientations of the two tryptophan radicals (W_C_ in RP_C_ and W_D_ in RP_D_).

**Figure 2.**
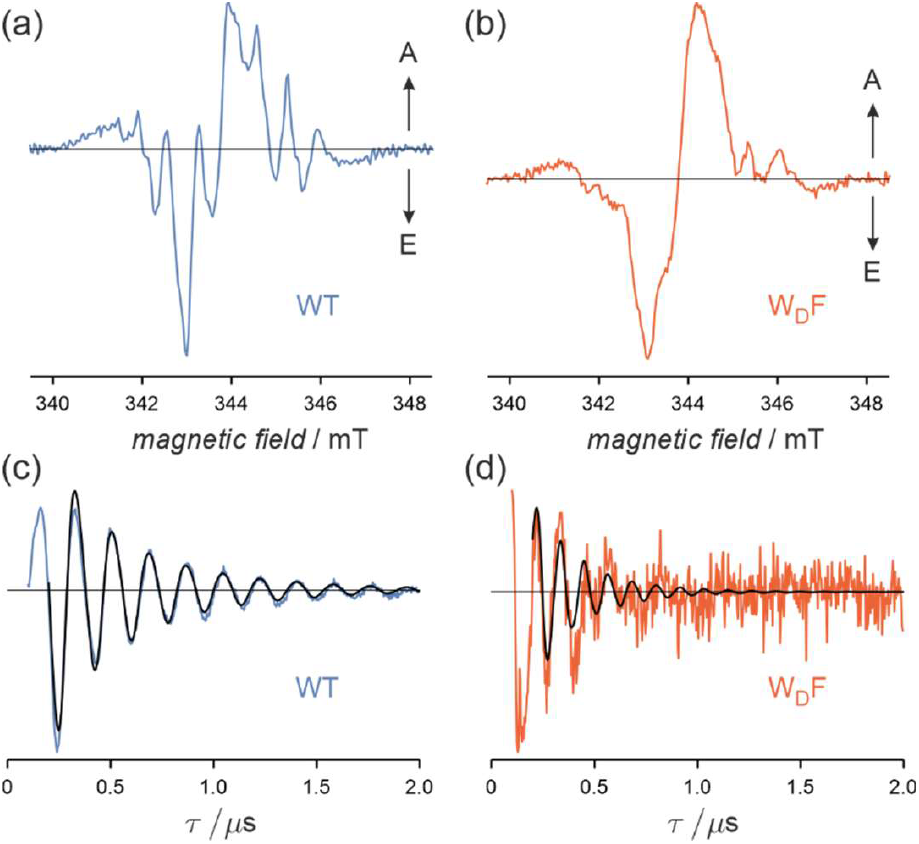
Time-resolved EPR spectra (a,b) and OOP-ESEEM spectra (c,d) of WT (a,c) and W_D_F (b,d) *Cj*CRY4a. The black lines in (c,d) are fits to Equation (1) using the parameters in Table 3. All spectra were acquired in the presence of 5 mM K_3_Fe(CN)_6_ to promote flavin re-oxidation.

Confirmation of the identities of the radical pairs is provided by out-of-phase electron spin echo envelope modulation (OOP-ESEEM) spectroscopy, an EPR technique ideally suited to measuring radical-radical separations in spin-correlated radical pairs [2, 54, 73]. Figures 2c,d show 34 GHz (Q-band) OOP-ESEEM data for WT and W_D_F *Cj*CRY4a recorded at 80 K with the first microwave pulse timed to come 200 ns after a 5 ns, 450 nm laser flash. Despite the much lower signal-to-noise ratio for the W_D_F mutant, it is clear that its modulation frequency exceeds that of the WT protein, reflecting the smaller separation of the radicals in RP_C_ compared to RP_D_.

**Table 3.** Parameters obtained from fitting the OOP-ESEEM data in Figure 2c,d.

| | $D$ / MHz | $J$ / MHz | $r$ / nm |
| --- | --- | --- | --- |
| WT *CjCRY4a* | $-8.54 \pm 0.09$ | $0.04 \pm 0.03$ | $2.09 \pm 0.01$ |
| W<sub>D</sub>F *CjCRY4a* | $-13.8 \pm 0.5$ | $0.29 \pm 0.17$ | $1.78 \pm 0.02$ |

Simulation of the data using Equation (1), based on a simple model [63] of the echo modulation, gave the results summarized in Table 3. The separations of the FAD^•−^ and TrpH^•+^ radicals (2.09 nm for the WT, 1.78 nm for W_D_F) match those reported for *Er*CRY4a and *Gg*CRY4a [2, 14], and are consistent with the FAD-Trp_D_ and FAD-Trp_C_ centre-to-centre distances, respectively, in the X-ray structure of *Cl*CRY4a [59] and homology models of *Er*CRY4a [2, 51] and *Gg*CRY4a [14].

### 3.3. Transient absorption spectroscopy

To determine the short-timescale kinetics of the photo-induced radicals, transient absorption spectra (Δ*A*, 400-800 nm) were recorded at 278 K as a function of the delay between a 15 ps, 450 nm pump pulse and a 1 ns, broadband (350-900 nm) probe pulse. Spectra are shown in Figures 3a,b for both forms of *Cj*CRY4a for pump-probe delays up to ∼1 μs. Negative signals between 400 and 500 nm arise from depletion of the ground state (FAD_OX_), whereas positive signals correspond to radicals formed by photo-induced electron transfer: FAD^•−^ (< 400 nm and 500-550 nm), Trp^•^ (500-550 nm), and TrpH^•+^ (500-620 nm). Similar spectra have been reported for *Gg*CRY4a [14].

**Figure 3.**
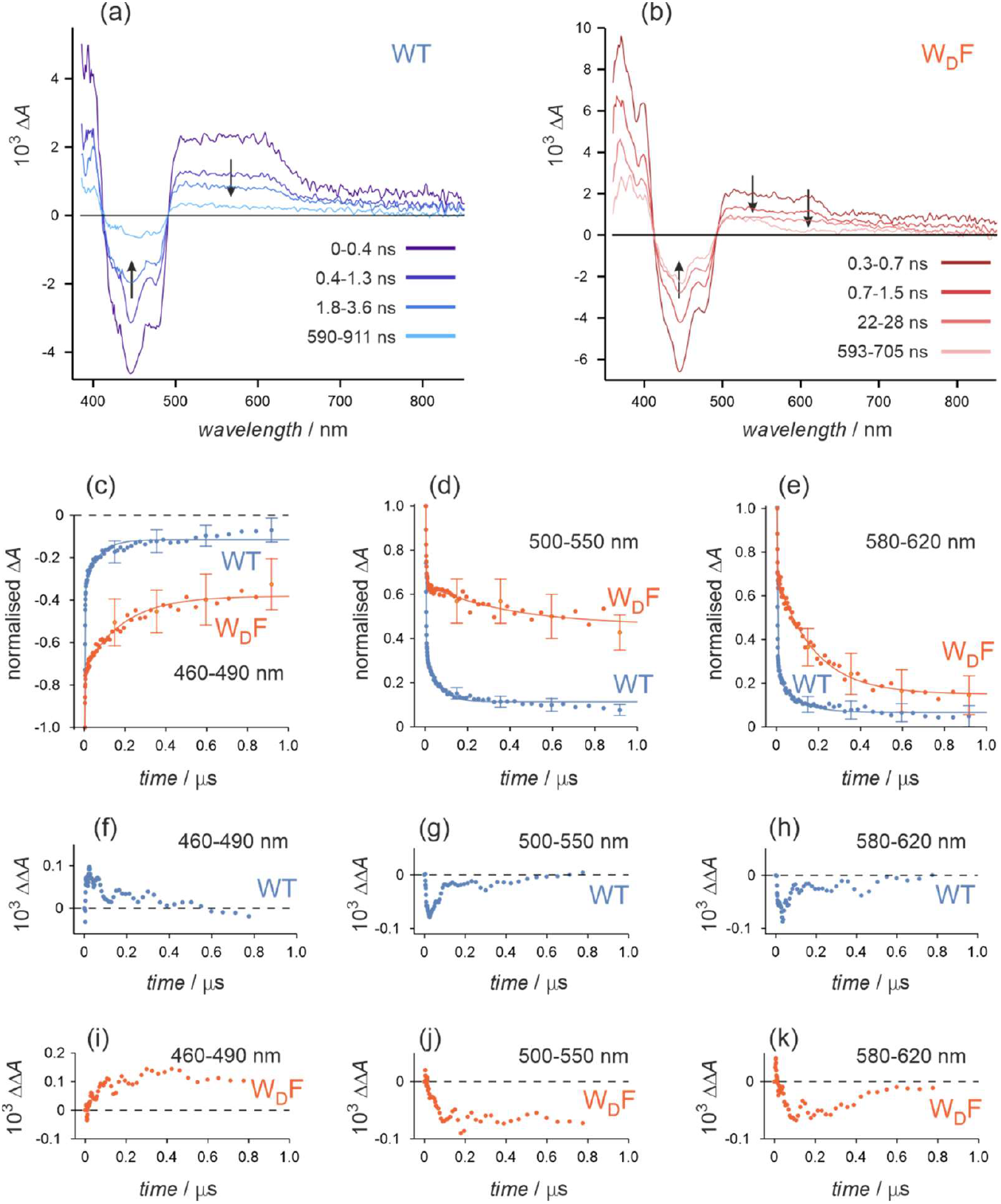
Transient absorption data for WT (blue) and W_D_F (red) forms of *Cj*CRY4a. (a,b) Spectra (Δ*A*) for different pump-probe delays. (c-e) Time-dependence of Δ*A* signals averaged over different wavelength bands. The lines are fits to a bi-exponential model (Supplementary Table S1). (f-k) Time dependence of ΔΔ*A* = Δ*A*(25 mT) − Δ*A*(0) signals. All data in (f-k) were smoothed using a 5-point moving average with 1:4:6:4:1 relative weights.

The major difference between WT and W_D_F *Cj*CRY4a occurs in the wavelength range 500-650 nm. The transient absorption signal of the WT protein in this region decays uniformly with increasing pump-probe delay, while the absorption of W_D_F between 550 and 650 nm decays more rapidly than between 500 and 550 nm absorption. From this, it is evident that in W_D_F the TrpH^•+^ radical deprotonates to form Trp^•^ on a < 1 μs timescale while in the WT protein, TrpH^•+^ remains protonated for more than 1 μs. Only by reducing the temperature to 268 K and increasing the glycerol content of the solution from 20% to 50% was it possible to see formation of Trp^•^ in WT *Cj*CRY4a on a sub-microsecond timescale (Supplementary Figure S3).

Unfortunately, it proved impossible to study WT and W_D_F *Cj*CRY4a using the same oxidant to convert the FAD radicals back to the ground state, FAD_OX_. Oxidation of FAD^•−^ in the WT protein by dissolved oxygen was inefficient and the W_D_F protein aggregated so extensively in the presence of ferricyanide as an oxidant that no useful data could be obtained. Measurements were therefore made using 1 mM Fe(III) for WT *Cj*CRY4a and air-equilibrated solutions for the W_D_F mutant. The difference in solution conditions limits the extent to which data for the two proteins can be directly compared.

Figures 3c-f show the time-dependence of the absorption signals of the two proteins in three wavelength bands. In all cases, the radical states decayed and the ground state recovered more rapidly for the WT than for the mutant, as found previously for *Gg*CRY4a [14]. Kinetic parameters obtained from bi-exponential fitting are given in Supplementary Table S1.

Together with the EPR results, these measurements demonstrate that the electron transfer chain and the radicals formed in response to photo-excitation of FAD in quail CRY4a are the same as in *Er*CRY4a and *Gg*CRY4a. This brings us to the essential question: is the observed photochemistry magnetically sensitive?

Figures 3g-j (WT) and 3k-n (W_D_F) show the differences between the transient absorption signals measured with and without a 25 mT magnetic field. Both proteins show clear magnetic field effects: negative for the radicals and positive for the ground state, consistent with singlet-born radical pairs [74]. As found for robin and chicken CRY4a’s [2, 14], magnetic field effects (= ΔΔ *A* / Δ *A*) on the order of 10% were observed for quail CRY4a, with the effects generally being larger and longer lived for the mutant than for the WT.

### 3.4. Cavity ring-down spectroscopy

Cavity ring-down spectroscopy (CRDS) achieves substantial improvements in detection sensitivity by exploiting the increased optical path lengths obtained by multiple passes of the probe light through a sample placed within an optical cavity [7, 64]. The price paid for this advantage is a reduced time-resolution (∼100 ns) compared to transient absorption. Figures 4a,b show the time-dependence of the CRDS signals, Δ*A*, recorded with and without a 30 mT applied magnetic field, whereas Figures 4c,d give the corresponding field-on minus field-off difference (MFE) signals, ΔΔ*A*, at 532 nm.

**Figure 4.**
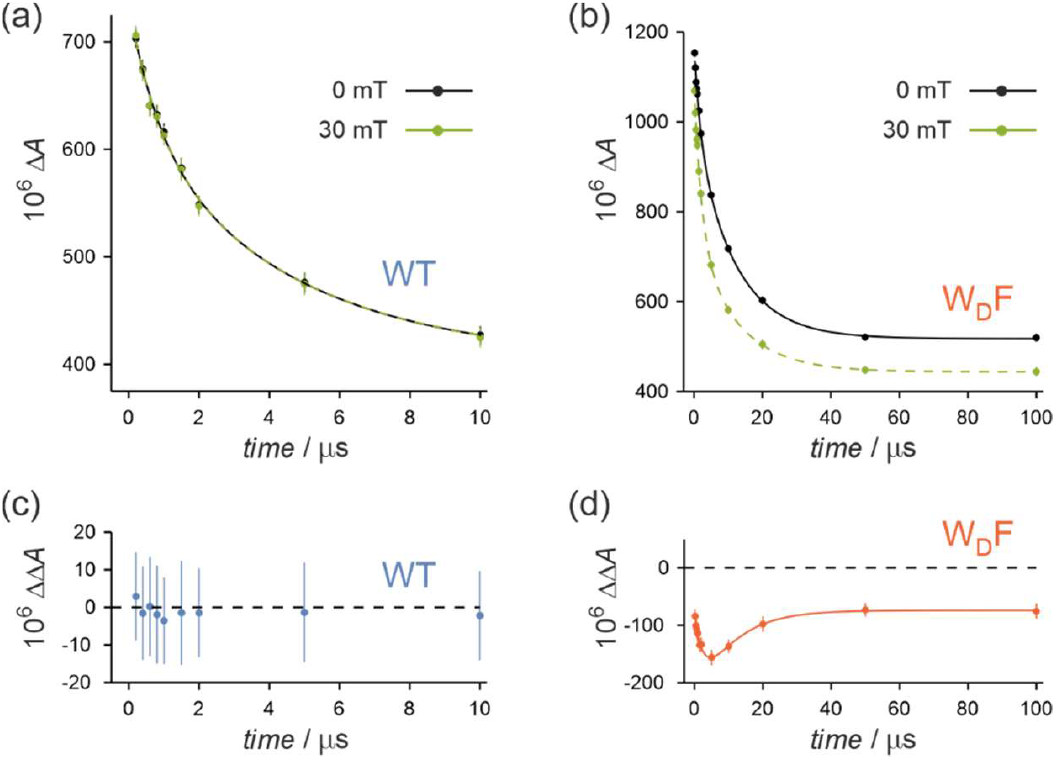
CRDS signals: Δ*A* (a,b) and ΔΔ*A* (c,d) of WT (a,c) and W_D_F (b,d) *Cj*CRY4a. The solid lines are fits using the parameters in Supplementary Table S2. All data were acquired in the presence of 5 mM K_3_Fe(CN)_6_ to promote flavin re-oxidation. Unlike transient absorption (Figure 3), these measurements did not suffer from protein aggregation in the presence of Fe(III). We attribute this difference to a combination of protein batch variability and differences in experimental conditions. Sample degradation is less pronounced in CRDS because of the lower concentrations, less extensive signal averaging, and smaller number of photons required to record the data

In agreement with the transient absorption measurements above, in which the magnetic field effects were seen to persist for approximately 100 ns (Figures 3f-k), no magnetic field effect could be observed for the WT protein (Figures 4a,c). W_D_F *Cj*CRY4a, by contrast, showed a higher transient yield of radicals and a large magnetic field effect (ca. −16% at 100 μs). Once again, the negative magnetic field effect on the radical concentration indicates a singlet-born radical pair.

### 3.5. Broadband cavity-enhanced absorption spectroscopy

The broadband cavity-enhanced absorption (BBCEAS) technique offers similar enhancements in detection sensitivity as well as broadband spectral resolution at the cost of a further reduction in time-resolution [45, 65]. Using continuous illumination (450 nm), absorption spectra can be obtained for transient intermediates that are long-enough lived to photo-accumulate. Figure 5a shows the difference between the Δ*A* signals measured with and without a 30 mT magnetic field, averaged over a period of 10 s. As above, the magnetic field effect, here observed as the change in the absorption of the protonated flavin radicals, FADH^•^, and of the deprotonated tryptophan radicals, Trp^•^, is stronger for W_D_F *Cj*CRY4a than it is for the WT protein. The quantum yields and time-evolution of the photo-induced radicals were broadly similar for the two proteins (Supplementary Figure S4).

**Figure 5.**
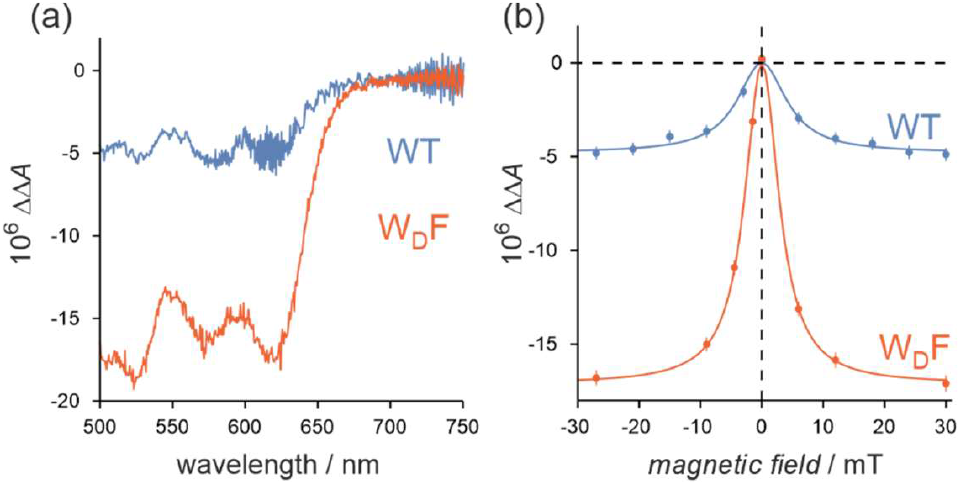
Spectrally resolved magnetic field effects in *Cj*CRY4a. BBCEAS ΔΔ*A* data for WT (blue) and W_D_F (red) *Cj*CRY4a. (a) Wavelength dependence. (b) Magnetic field dependence. The solid lines in (b) are fits to a Lorentzian model. All data were acquired for samples saturated with O_2_ (1 atm) to promote flavin re-oxidation.

The sensitivity afforded by BBCEAS allows changes in absorption signals to be measured as a function of magnetic field strength. Data for the two forms of *Cj*CRY4a are shown in Figure 5b. The shapes of these plots are characteristic of radical pairs with ∼1 mT hyperfine interactions and have similar values of *B*_1/2_, the magnetic field strength at which ΔΔ*A* equals the mean of its value at zero field and the asymptotic value at high field: 4.8 ± 1.9 mT for the WT and 3.3 ± 0.3 mT for W_D_F. These values are typical for [FAD^•−^ TrpH^•+^] radical pairs in cryptochromes [2, 14, 45].

### 3.6. Fluorescence microscopy

Detection of the fluorescence of FAD in cryptochromes can be used as an alternative to optical absorption spectroscopy [66-68]. Figure 6 shows confocal fluorescence microscopy data for the two forms of *Cj*CRY4a. The protein solutions were irradiated continuously at 448 nm while a 17 mT magnetic field was repeatedly switched on and off every 95.9 s. The fluorescence of the excited singlet state of FAD_OX_ rises when the field is applied and falls when it is removed because the magnetic field changes the probability of forming long-lived FADH^•^ and hence, also, the steady state concentration of FAD_OX_. Because the fluorescence monitors FAD_OX_ rather than the radicals, the magnetic field effect is positive, as expected for singlet-born radical pairs [67, 75].

**Figure 6.**
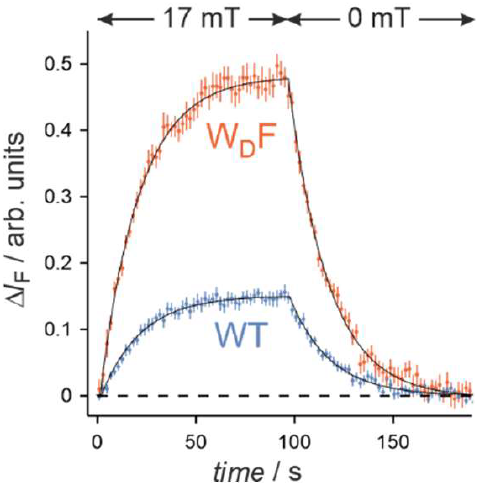
Time-dependence of the change in the fluorescence intensity (Δ*I*_F_) of WT (blue) and W_D_F (red) *Cj*CRY4a resulting from switching a 17 mT magnetic field on and off. The lines are single-exponential fits of the signal rise (field on) and fall (field off), with time constants 21.2 ± 0.6 s (WT) and 20.1 ± 0.3 s (W_D_F). All data were acquired for samples saturated with O_2_ (1 atm) to promote flavin re-oxidation.

Apart from a three-fold difference in the magnetic field effect (again, W_D_F > WT), the two proteins have very similar behaviour with ca. 20 s rise and fall times on field switching.

## 4. Discussion

Our study demonstrates that *Cj*CRY4a has similar magnetic reactivity to *Er*CRY4a and *Gg*CRY4a. One could expect the photochemistry of quail CRY4a to resemble that of the robin and chicken proteins, considering that all three CRY4a proteins have high sequence homology in the neighbourhood of the FAD/Trp electron transfer chain. The only differences occur near W_C_ and W_D_ at positions 317 and 320 where *Er*CRY4a has a cysteine and a lysine, respectively, whereas *Gg*CRY4a and *Cj*CRY4a both have an arginine and a glutamic acid, respectively. Despite the different charges of the side chains, the *Gg*CRY4a mutants R317C and E320K show similar magnetic field effects to WT *Gg*CRY4a and WT *Er*CRY4a [14]. With our current knowledge and experimental conditions, differences in the abilities of (migratory) robins and (non-migratory) chickens to orient in the Earth’s magnetic field cannot be ascribed to these sequence variations [14].

*Cj*CRY4a binds FAD and forms FAD^•−^-TrpH^•+^ radical pairs when excited by blue light. For the WT protein, with an intact Trp-tetrad, four sequential electron transfers along the Trp-tetrad result in a radical pair comprising the radical forms of the flavin and the terminal tryptophan, W_D_ (Figure 1). When the tetrad is truncated by replacing this residue with a phenylalanine, the photo-induced radical pair instead contains the radical form of W_C_. The subsequent reactions of both radical pairs — RP_D_ in the WT and RP_C_ in the W_D_F mutant — are affected by weak applied magnetic fields, with the W_D_F mutant considerably more sensitive than the WT (Figures 3-6). All of these observations mirror the behaviour of *Gg*CRY4a and *Er*CRY4a [2, 14].

The different magnetic sensitivities of the WT and W_D_F quail proteins are interesting not least because one might have predicted that the fourth tryptophan, present in avian CRY4as but not in the cryptochromes of organisms that seem to have much less of a need of a magnetic compass, e.g. plants, could result in larger magnetic field effects than possible with a Trp-triad in, for example, *Arabidopsis thaliana* CRY1 [74, 76, 77]. As in the case of *Gg*CRY4a, the smaller magnetic field effects for the WT protein can be traced to the relative rate constants for TrpH^•+^ deprotonation (*k*_dep_) and FAD^•−^ TrpH^•+^ recombination (*k*_rec_) which were here found to be in the order (Figures 3 and 4):

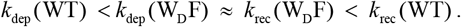

>Other things being equal, a strong magnetic field effect requires *k*_dep_ and *k*_rec_ to be similar [2, 74]. This is evidently the case for the W_D_F mutant but not for the WT. That radical recombination should be slower for RP_C_ in W_D_F than for RP_D_ in the WT protein, seems counterintuitive given the larger separation of the radicals in the latter. One possibility is that the back electron transfer rates are dominated by differences in the thermodynamic driving force and/or the reorganisation energy [14]. In relation to the smaller magnetic field effect in the WT protein, we have speculated that the fourth tryptophan in avian CRY4a may have allowed evolution to optimise the sensing and signalling functions of a cryptochrome magnetoreceptor in a way that would not have been possible with only a Trp-triad [2, 3].

The present findings that *Cj*CRY4a has magnetic properties similar to robin CRY4a underline the validity of potentially using domesticated quails as a model animal in magnetoreception and orientation experiments. However, if quail were to be used for behaviour experiments, hybrids with wild birds would have to be created to regain the *Zugunruhe* phenotype, which is much reduced or absent in pure domestic *C. japonica* [41]. Irrespective of its phenotype restoration, hybrids are essential because quail are generally susceptible to inbreeding. The amino acid sequences of the two CRY4a from wild *C. coturnix* and *C. japonica* are essentially identical, and four recurrent variants (H4R, A20S, R462Q, and A529T) are found in both wild quail species (Table 1). These natural recurrent variants are all located away from FAD and the Trp tetrad, based on the crystal structure of *Cl*CRY4, and in addition, the effects of these missense variants were predicted to be benign (Table 2). This suggests that any wild, domesticated, or hybrid quail involving any combination of *C. coturnix* and *C. japonica* will have a magnetosensitive CRY4a.

While captive breeding enables precise control over environmental conditions, it is extremely challenging to provide captive-bred birds with all the natural pre-migratory inputs and to facilitate the natural learning processes they go through in the wild. For instance, pre-migratory flights would certainly not be possible. Such flights are thought to be used to practice flight skills, to assess meteorological conditions aloft, and/or to create magnetic, olfactory, or visual landscape maps for navigation [78-80]. For this reason and because of their distinct phylogenetics, it is therefore unlikely that quails can completely replace wild-caught night-migratory passerines to study the complex navigation system. However, establishing an additional model species that can be bred in captivity could be beneficial for studying different aspects of radical-pair-based magnetoreception, including electrophysiology, retinal circuitry, or neurobiological studies. The generation of knock-out and knock-in quails using genome editing [81] and pharmacological interventions, combined with electrophysiological and behavioural analyses, would advance research on magnetoreception, orientation, and navigation.

In conclusion, we have confirmed that the magnetic responses of quail CRY4a are similar to those of robin CRY4a, supporting the idea that quail could be a promising additional experimental model for investigating the intricacies of CRY4a-mediated magnetoreception.

## Acknowledgements

We are grateful to Lisa Borowski, Simon Horst, Stefanie Käsehagen, Katarina Karm, Hannah Käse, Wiemke Reiners, and Thore van Düllen and for helping with protein expression and purification. We thank the animal facilities at the University of Oldenburg for excellent animal keeping and care taking. We are grateful to the Oxford Chemistry buildings and maintenance staff for their recent assistance in moving the magnetic field effect laboratories to the Chemistry Research Laboratory.

## Author contributions

HM, SRM, CRT, PJH, and RB conceived and supervised the study. RB, MB, and JS cloned and established expression and purification protocols for *Cj*Cry4a WT and *Cj*CRY4a W369F. RB and JS expressed and purified or supervised the expression and purification of all the protein samples, with help from GD, GS, JJF, MK, SA, and JX. TK did the sequence analyses. All spectroscopic data were recorded in Oxford. TLP performed the EPR experiments and analysed the data. KBH and HH conducted the TA experiments, the results of which were analysed by KBH and JG. DRC undertook the CRDS measurements and analysed the data. PDFM, DRC and CO performed the BBCEAS measurements, with PDFM conducting the analysis. JG and LMA performed the confocal microscopy experiments, with JG undertaking the analysis. OP-L assisted with the spectroscopic measurements. RB and PJH wrote the initial draft of the manuscript, which was refined by HM, TK, CRT and SRM. All authors contributed to the interpretation of the data and commented on the manuscript.

## Ethics

This work did not require ethical approval from a human subject or animal welfare committee. The permission to keep and collect tissues from quails was provided by Niedersächsisches Landesamt für Verbraucherschutz und Lebensmittelsicherheit (LAVES).

## Declaration of AI use

We have not used AI-assisted technologies in creating this article.

