## Supplements for "Magnetic sensitivity of cryptochrome 4a in domesticated quail with migratory origins"

### Supplementary Materials

#### S1. Sequence alignments

```
ErCry4a      MLHRTIHLFRKELRLHDNPVLLAALQSSEALYPVYILDRAFLTSSMHIGALRWHFLLQSL    60
GgCry4a      MRHRTIHLFRKGLRLHDNPALLAALQSSEVVYPVYILDRAFMTSSMHIGALRWHFLLQSL    60
CjCry4a      MRHHTIHLFRKGLRLHDNPALLAALQSSEVVYPVYILDRAFMTSVMHIGALRWHFLLQSL    60
CcCry4a      MRHRTIHLFRKGLRLHDNPALLAALQSSEVVYPVYILDRAFMTSVMHIGALRWHFLLQSL    60
             * *:******* ******* *********.:**********:** ***************

ErCry4a      EDLHKNLCQLGSCLLVIQGEYETVLRDHIQKWSITQVTLDAEMEPFYKEMEANIQCLGAE    120
GgCry4a      EDLRSSLRQLGSCLLVIQGEYESVVRDHVQKWNITQVTLDAEMEPFYKEMEANIRGLGEE    120
CjCry4a      EDLRSSLCQLGSCLLVIQGEYESVVRDHVQKWNITQVTLDEEMEPFYKEMEANIRGLGEE    120
CcCry4a      EDLRSSLCQLGSCLLVIQGEYESVVRDHVQKWNITQVTLDEEMEPFYKEMEANIRGLGEE    120
             ***:..* **************:*:***:***.******* *************: ** *

ErCry4a      LGFEVLSLGSHSLYDTQRILDINGGSPPLTYKRFLHILSLLGDPEVPVRNLTAEDFQRCS    180
GgCry4a      LGFQVLSLMGHSLYNTQRILELNGGTPPLTYKRFLRILSLLGDPEVPVRNPTAEDFQRCS    180
CjCry4a      LGFEVLSLMGHSLYNTQRILELNGGTPPLTYKRFLRILALLGDPEVPVRNLTAEDFQRCS    180
CcCry4a      LGFEVLSLMGHSLYNTQRILELNGGTPPLTYKRFLRILALLGDPEVPVRNLTAEDFQRCS    180
             ***:**** .****:*****::***:*********:**:*********** *********

ErCry4a      APDPDLAECYRVPLPVDLKISPENLSPWRGGETEGLQRLEQHLTDQGWVASFTKPRTIPN    240
GgCry4a      PPELGLAECYGVPLPTDLKIPPESISPWRGGESEGLQRLEQHLADQGWVASFTKPKTVPN    240
CjCry4a      PPELGLAERYGVPLPTDLKIPPESISPWRGGESEGLHRLEQHLADQGWVASFTKPKTIPN    240
CcCry4a      PPELGLAERYGVPLPTDLKIPPESISPWRGGESEGLHRLEQHLADQGWVASFTKPKTIPN    240
              *: .*** * ****.**** **.:*******:***:******:***********:*:**

ErCry4a      SLLPSTTGLSPYFSMGCLSVRTFFYRLSNIYAQAKHHSLPPVSLQGQLLWREFFYTVASA    300
GgCry4a      SLLPSTTGLSPYFSTGCLSVRSFFYRLSNIYAQAKHHSLPPVSLQGQLLWREFFYTVASA    300
CjCry4a      SLLPSTTGLSPYFSMGCLSVRSFFYRLSNIYAQAKHHSLPPVSLQGQLLWREFFYTVASA    300
CcCry4a      SLLPSTTGLSPYFSMGCLSVRSFFYRLSNIYAQAKHHSLPPVSLQGQLLWREFFYTVASA    300
             ************** ******:**************************************

ErCry4a      TPNFTQMAGNPICLQICWYKDAERLHKWKMAQTGFPWIDAIMTQLRQEGWIHHLARHAVA    360
GgCry4a      TPNFTKMAGNPICLQIRWYEDAERLHKWKTAQTGFPWIDAIMTQLRQEGWIHHLARHAAA    360
CjCry4a      TPNFTKMAGNPICLQIRWYEDAERLHRWKTAQTGFPWIDAIMTQLRQEGWIHHLARHAAA    360
CcCry4a      TPNFTKMAGNPICLQIRWYEDAERLHRWKTAQTGFPWIDAIMTQLRQEGWIHHLARHAAA    360
             *****:********** **:******:** ****************************.*

ErCry4a      CFLTRGDLWISWEEGMKVFEELLLDADYSINAGNWMWLSASAFFHQYTRIFCPVRFGKRT    420
GgCry4a      CFLTRGDLWISWEEGMKVFEELLLDADYSINAGNWMWLSASAFFHHYTRIFCPVRFGRRT    420
CjCry4a      CFLTRGDLWISWEEGMKVFEELLLDADYSINAGNWMWLSASAFFHQYTRIFCPVRFGRRT    420
CcCry4a      CFLTRGDLWISWEEGMKVFEELLLDADYSINAGNWMWLSASAFFHQYTRIFCPVRFGRRT    420
             *********************************************:***********:**

ErCry4a      DPQGNYIRKYLPILKNFPSKYIYEPWTASEEEQKQAGCIIGRDYPFPMVNHKEASDHNLQ    480
GgCry4a      DPEGQYIRKYLPILKNFPSKYIYEPWTASEEEQKQAGCIIGRDYPFPMVDHKEASDHNLQ    480
CjCry4a      DPEGEYIRKYLPILKNFPSKYIYEPWTASEEEQKQAGCIIGQDYPFPMVDHKEASDHNLQ    480
CcCry4a      DPEGEYIRKYLPILKNFPSKYIYEPWTASEEEQKQAGCIIGQDYPFPMVDHKEASDHNLQ    480
             **:*:************************************:*******:**********

ErCry4a      LMRQVREEQHRTAQLTRDDADDPMEIKVKRDHTEENISKGKVARTTE--      527
GgCry4a      LMKQAREEQHRIAQLTRDDADDPMEMKLKRDHSEESFTKTKAARMTEQT      529
CjCry4a      LMKQVREEQYRTAQLTRDDADDPMEMKLKRDLSEENLPKTKAARMTEQA      529
CcCry4a      LMKQVREEQYRTAQLTRDDADDPMEMKLKRDLSEENLPKTKAARMTEQA      529
             **:*.****:* *************:*:*** :**.: * *.** **
```

**Figure S1**. Sequence alignments comparing chicken (*Gg*), quail (*Cj* and *Cc*) and robin (*Er*) WT CRY4a (Clustal Omega (RRID:SCR_001591)). *Cj*CRY4a, used in this study, differs only in one amino acid (position 4) from *Cc*CRY4a (*C. coturnix* SAMN19229867, collected in Spain, Table 1). *Cj*CRY4a is 96% identical to *Gg*CRY4a, and 88% identical to *Er*CRY4a [82]. The four tryptophans in the Trp-tetrad are highlighted in green. The amino acid residues at positions 317 and 320 that are in close proximity to the electron transfer chain are highlighted in yellow. Positions of natural missense variants leading to four amino acid polymorphisms in both *Cc*CRY4a and *Cj*CRY4a are highlighted in turquoise.

#### S2. UV-vis spectra

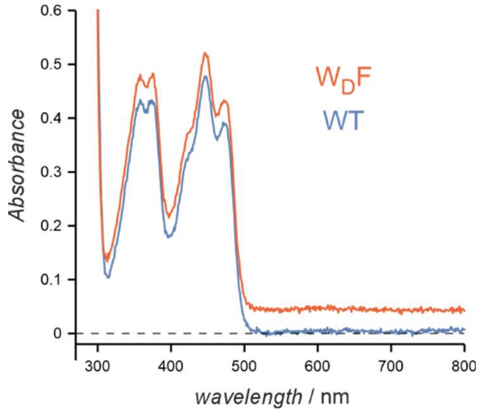


**Figure S2**. UV-visible spectra of WT *Cj*CRY4a and its W$_D$F mutant showing that the FAD is correctly bound and in its fully oxidised state, FAD$_{OX}$. The spectrum of W$_D$F has been offset by 0.05 Absorbance units for clarity.

#### S3. Deprotonation of tryptophan radical in WT *Cj*CRY4a

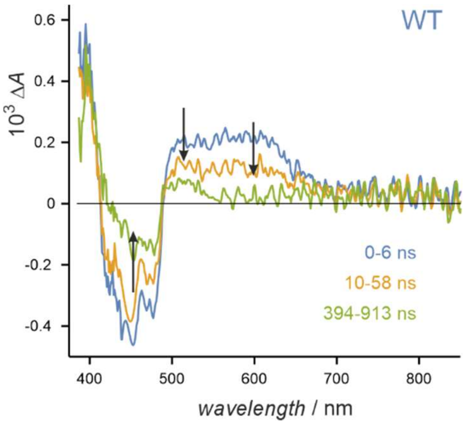


**Figure S3**. Transient absorption data for WT *Cj*CRY4a at 268 K in 50% glycerol in water. The faster decay at ~600 nm than at ~520 nm results from the conversion of $Trp_DH^{\bullet+}$ to $Trp_D^{\bullet}$.

#### S4. Analysis of transient absorption data for *Cj*CRY4a

TA provides information on the complex kinetics of the production of radicals, in particular $FAD^{•-}$, $TrpH^{•+}$, and $Trp^{•}$ and their decay back to the ground state. The rate of $FAD^{•-}$ $TrpH^{•+}$ recombination in particular, depends strongly on the distance between the radicals and hence on the extent of the electron transfer down the tryptophan tetrad/triad. Back electron transfer $FAD^{•-}$ to $Trp_AH^{•+}$ occurs on a (sub-)ps timescale, *i.e.*, too fast to observe with this method. The TA kinetics clearly show multiphasic behaviour. For simplicity, we fit to a biexponential decay of the form,

$$A(t) = A_1 \exp(-t/\tau_1) + A_2 \exp(-t/\tau_2) + A_3 .$$

We interpret the faster of the two decays ($\tau \approx 1$ ns) as arising from back electron transfer from $FAD^{•-}$ to $Trp_BH^{•+}$. The slower decay probably includes a range of processes including recombination of $FAD^{•-}$ with $Trp_CH^{•+}$ (and $Trp_DH^{•+}$) as well as radical protonation/deprotonation kinetics.

| **WT** | $10^3 A_1$ | $\tau_1$ / ns | $10^3 A_2$ | $\tau_2$ / ns | $10^3 A_3$ |
|---|---|---|---|---|---|
| 460-490 nm | –0.67 | 1.2 ± 0.5 | –0.21 | 68 ± 7 | –0.12 |
| 500-550 nm | 0.65 | 1.1 ± 0.5 | 0.24 | 55 ± 4 | 0.11 |
| 580-620 nm | 0.69 | 1.0 ± 0.5 | 0.24 | 52 ± 4 | 0.07 |

| **W$_D$F** | $10^3 A_1$ | $\tau_1$ / ns | $10^3 A_2$ | $\tau_2$ / ns | $10^3 A_3$ |
|---|---|---|---|---|---|
| 460-490 nm | –0.25 | 1.3 ± 0.5 | –0.36 | 188 ± 14 | –0.39 |
| 500-550 nm | 0.35 | 1.6 ± 0.5 | 0.18 | 376 ± 80 | 0.47 |
| 580-620 nm | 0.34 | 0.9 ± 0.5 | 0.52 | 181 ± 8 | 0.14 |

**Table S1**. Parameters obtained by fitting the data shown in Figure 3c-f to the biexponential model function $A(t) = A_1 \exp(-t/\tau_1) + A_2 \exp(-t/\tau_2) + A_3$.

#### S5. Analysis of CRDS data for *Cj*CRY4a

| **WT** | $10^4 A_1$ | $\tau_1$ / µs | $10^4 A_2$ | $\tau_2$ / µs | $10^4 A_3$ |
|---|---|---|---|---|---|
| $\Delta A(B = 0)$ | 1.17 ± 0.06 | 2.01 ± 0.71 | 0.28 ± 0.07 | 20.99 ± 0.25 | 1.62 ± 0.04 |
| $\Delta A(B = 30\text{ mT})$ | 1.21 ± 0.11 | 1.95 ± 0.19 | 0.26 ± 0.08 | 24.04 ± 1.76 | 1.61 ± 0.06 |
| **W$_D$F** | | | | | |
| $\Delta A(B = 0)$ | 4.17 ± 0.06 | 12.35 ± 0.71 | 2.13 ± 0.07 | 2.57 ± 0.25 | 4.99 ± 0.04 |
| $\Delta A(B = 30\text{ mT})$ | 3.47 ± 0.11 | 2.35 ± 0.19 | 2.84 ± 0.08 | 14.59 ± 1.76 | 4.24 ± 0.06 |
| $\Delta\Delta A$ | –1.50 ± 0.07 | 8.27 ± 1.06 | 1.50 ± 0.10 | 2.17 ± 0.35 | –0.73 ± 0.05 |

**Table S2**. Parameters obtained by fitting the data shown in Figure 4a,b,d to the biexponential model function $A(t) = A_1 \exp(-t/\tau_1) + A_2 \exp(-t/\tau_2) + A_3$.

#### S6. BBCEAS data for *Cj*CRY4a

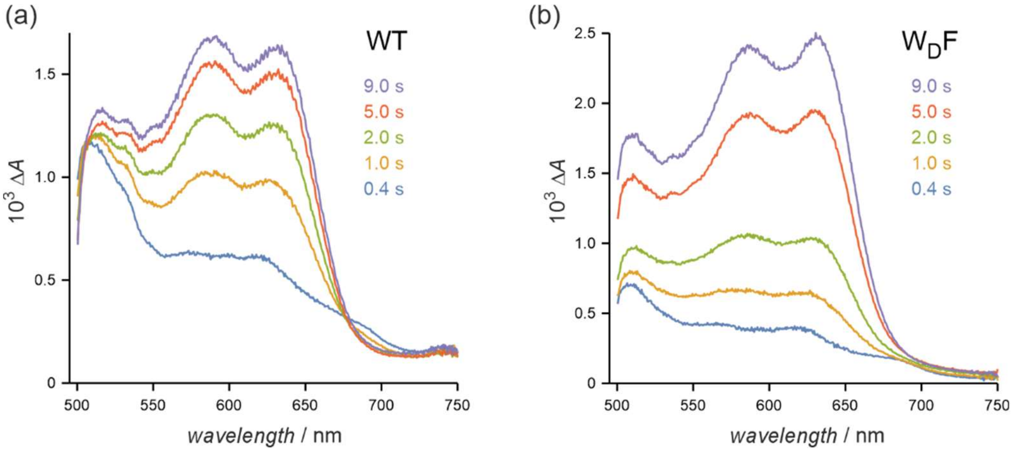


**Figure S4**. BBCEAS $\Delta A$ data for (a) WT and (b) $W_DF$ *Cj*CRY4a as a function of total illumination time.
